# An Aryl Hydrocarbon Receptor from the Caecilian *Gymnopis multiplicata* Suggests Low Dioxin Affinity in the Ancestor of All Three Amphibian Orders

**DOI:** 10.1101/750653

**Authors:** Sarah A. Kazzaz, Sara Giani Tagliabue, Diana G. Franks, Michael S. Denison, Mark E. Hahn, Laura Bonati, Wade H. Powell

## Abstract

The aryl hydrocarbon receptor (AHR) plays pleiotropic roles in the development and physiology of vertebrates in conjunction with xenobiotic and endogenous ligands. It is best known for mediating the toxic effects of dioxin-like pollutants such as 2,3,7,8-tetracholordibenzo-*p*-dioxin (TCDD). While most vertebrates possess at least one AHR that binds TCDD tightly, amphibian AHRs bind TCDD with very low affinity. Previous analyses of AHRs from *Xenopus laevis* (a frog; order Anura) and *Ambystoma mexicanum* (a salamander; order Urodela) identified three amino acid residues in the ligand-binding domain (LBD) that underlie low-affinity binding. In *X. laevis* AHR1β, these are A354, A370, and N325. Here we extend the analysis of amphibian AHRs to the caecilian *Gymnopis multiplicata*, representing the remaining extant amphibian order, Apoda. *G. multiplicata* AHR groups with the monophyletic vertebrate AHR/AHR1 clade. The LBD includes all three signature residues of low TCDD affinity, and a structural homology model suggests that its architecture closely resembles those of other amphibians. In transactivation assays, the EC50 for reporter gene induction by TCDD was 17.17 nM, comparable to *X. laevis* AhR1β (26.23 nM) and *Ambystoma* AHR (34.09 nM) and dramatically higher than mouse AhR (0.13 nM), a trend generally reflected in direct measures of TCDD binding. These shared properties distinguish amphibian AHRs from the high-affinity proteins typical of both more ancient vertebrate groups (teleost fish) and those that appeared more recently (tetrapods). We suggest that AHRs with low TCDD affinity represent a basal characteristic that evolved in a common ancestor of all three extant amphibian groups.

**Research Highlights:**

- A caecilian aryl hydrocarbon receptor exhibits low dioxin binding and sensitivity.
- The protein’s ligand-binding domain resembles frog and salamander AHRs in structure and function.
- AHR with low dioxin affinity likely evolved in a common ancestor of all three extant amphibian groups.

**Graphical Abstract:** 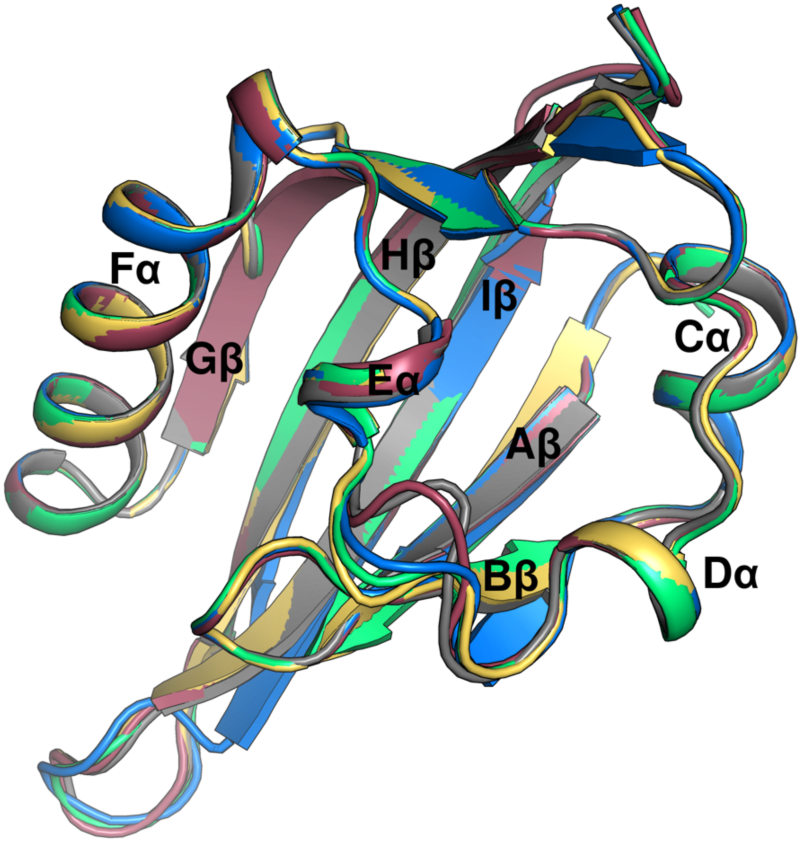

## Introduction

The aryl hydrocarbon receptor (AHR), a member of the Per-ARNT-Sim (bHLH/PAS) family of proteins (McIntosh, Hogenesch, & Bradfield, 2010), is a ligand-activated transcription factor that mediates the toxicity of a wide range of environmental contaminants, including chlorinated dioxin-like compounds and polynuclear aromatic hydrocarbons (Denison, Soshilov, He, DeGroot, & Zhao, 2011). The unliganded receptor is part of a cytoplasmic complex with hsp90, p23, and AIP (Murray & Perdew, 2011). Agonist binding triggers translocation to the nucleus, shedding of these chaperones, and dimerization with the ARNT protein. The AHR:ARNT heterodimer binds to cognate enhancer sequences (Seok et al., 2017) and alters transcription of numerous target genes (Frueh, Hayashibara, Brown, & Whitlock, 2001; Puga, Maier, & Medvedovic, 2000). Well-known gene targets include the “AHR gene battery,” encoding phase I and phase II detoxification enzymes such as the cytochrome P450 family 1 members (*CYP1*s) and UDP glucuronosyltransferase, glutathione S transferase, and quinone reductase (Nebert et al., 2000). Induced over two orders of magnitude, *CYP1*s are frequently used as a biomarker of exposure to AHR agonists (Hahn, 2002). AHR also exerts biological effects through “non-classical” mechanisms involving interactions with additional signaling pathways and nuclear proteins (Denison et al., 2011).

AHR can be activated by structurally diverse agonists of both xenobiotic and endogenous origin. The prototypical agonist in mechanistic toxicological studies of AHR is the industrial contaminant 2,3,7,8-tetracholorodibenzo-*p*-dioxin (TCDD, dioxin). Among the most toxic planar halogenated aromatic hydrocarbons, TCDD binds vertebrate AHRs with relatively high affinity (Van den Berg et al., 2006). Another potent AHR agonist, 6-formylindolo[3,2-*b*]carbazole (FICZ), has endogenous sources, forming *in vivo* from tryptophan as a photoproduct, an oxidation product, or as a metabolite produced by gut bacteria (Rannug & Rannug, 2018). FICZ binds AHR with even higher affinity than TCDD, but it is rapidly metabolized by the detoxification enzymes induced by exposure (Wincent et al., 2009).

Dioxin toxicity varies substantially between different vertebrate species or populations. Many studies link differences in AHR ligand binding affinity to these variations. Invertebrate AHRs typically do not bind TCDD, and the animals are largely insensitive to its toxic effects (Hahn, Karchner, & Merson, 2017). TCDD-sensitive mouse strains express the high-affinity *AHR^b-1^* allele, while toxicity is lower in strains expressing *AHR^d^*, which encodes a protein that binds TCDD with 4- to 10-fold lower affinity, comparable to the human receptor (Ema et al., 1994; Poland, Palen, & Glover, 1994; Ramadoss & Perdew, 2004). Affinity for AHR also underlies wide-ranging variations in TCDD toxicity between many bird species (Farmahin et al., 2013; Karchner, Franks, Kennedy, & Hahn, 2006). Amphibian receptors display less robust xenobiotic binding (Lavine et al., 2005; Shoots et al., 2015), with *X. laevis* AHRs exhibiting K_d_’s 20- to 50-fold higher than many fish, birds, and mammals (Karchner, Franks, & Hahn, 2005; Karchner et al., 2006) for TCDD. Amphibians are concomitantly much less sensitive to toxicity of dioxin-like chemicals (Beatty, Holscher, & Neal, 1976; Collier, Orr, Morris, & Blank, 2008; Jung & Walker, 1997; Pezdirc, Heath, Bizjak Mali, & Bulog, 2011; Vajda & Norris, 2005) and polynuclear aromatic hydrocarbons (Fort, James, & Bantle, 1989; Propst, Fort, Stover, Schrock, & Bantle, 1997).

To elucidate the structural basis for differences in TCDD affinity, ligand-binding domains (LBDs) have been characterized in AHRs from animals across the spectrum of TCDD sensitivity. A comparative study of AHR1 from the chicken (*Gallus gallus*) and common tern (*Sterna hirundo*) identified two residues—I324 and S380—that are critical for high-affinity binding by the chicken receptor, the species more sensitive to TCDD toxicity (Karchner et al., 2006). In mice, a single polymorphism (A375V) reduces TCDD affinity 4-5 fold in the AHR^d^ allele vs. AHR^b-1^; the LBD from human AHR contains valine at the homologous position (Ema et al., 1994; Poland et al., 1994; Ramadoss & Perdew, 2004). Additional signature residues conferring high-affinity binding in mouse AHR (Pandini et al., 2009) and AHRs from other species (Fraccalvieri et al., 2013) were identified by homology modeling and confirmed using site-directed mutagenesis. Similar approaches were employed to explain the low TCDD affinity of amphibian AHRs. Three amino acids within the LBD of *X. laevis* AHR1β confer low-affinity binding—N325, A354, and A370 (Odio et al., 2013). Homologous residues are present in AHR from *X. tropicalis* (Order Anura; frogs and toads) and *Ambystoma mexicanum* (Order Urodela; salamanders (Shoots et al., 2015). These shared sequence elements and functional properties suggest that low TCDD affinity emerged in the common ancestor of these two extant amphibian groups. Is it possible that low TCDD affinity appeared even earlier in amphibian evolution?

Caecilians, the legless amphibians, comprise Order Apoda, the earliest of the extant orders to diverge from the common lineage (Hay, Ruvinsky, Hedges, & Maxson, 1995; Pyron & Wiens, 2011; Zardoya & Meyer, 2001; Zhang & Wake, 2009; Zhang, Zhou, Chen, Liu, & Qu, 2005). Therefore, caecilians offer an opportunity to probe the timing of emergence of low affinity AHRs during amphibian evolution. If the caecilians possess a low-affinity AHR, then this phenotype is likely a basal feature, arising in an ancestor of all extant groups. Alternatively, if caecilian AHRs bind dioxin-like compounds with high affinity—similar to AHRs from both more ancient and more recent vertebrate groups— low-affinity binding must have evolved in the common ancestor of frogs and salamanders after the caecilian divergence. This study sought to address this question. In the first characterization of a caecilian AHR, we report the sequence, pharmacological properties, and a structural model of the ligand binding domain of a receptor derived from the Varagua caecilian, *Gymnopis multiplicata*.

## Materials and Methods

### AHR Ligands

TCDD was obtained from ULTRA Scientific dissolved in toluene. This material was dried under a N_2_ stream, dissolved in DMSO, and stored at room temperature. 6-formylindolo[3,2-*b*]carbazole (FICZ; Enzo Life Sciences; lyophilized powder) was dissolved in DMSO and stored in the dark at −20°C.

### cDNA cloning

Total RNA was extracted from an undescribed *Gymnopis multiplicata* tissue sample provided by the Museum of Vertebrate Zoology at the University of California, Berkeley. The sample (MVZ:Herp:269228) was collected in Nicaragua in 2010 and stored subsequently in RNAlater (Ambion). Total RNA was purified using RNA STAT-60 (Tel-Test Inc.). A partial cDNA encoding AHR was amplified by RT-PCR using the GeneAmp Gold RNA PCR Reagent Kit (Applied Biosystems). Total RNA (1 µg per reaction) was reverse transcribed with random hexamers as directed by the manufacturer. Degenerate PCR primers (Supplemental Information, Table S1; 1 µM each) were designed from regions of conserved amino acid sequence and have been used previously to amplify AHRs sequences from a variety of vertebrates, including teleost fish, frogs and salamanders, and birds (*e.g*. (Hahn & Karchner, 1995; Karchner et al., 2006; Lavine et al., 2005; Shoots et al., 2015). Thermocycler conditions were 95°C for 10 min; 43 cycles of 95°C for 15 seconds, 50°C for 30 sec, 72°C for 1 min; and 72°C for 7 min. Initial RT-PCR reactions were diluted 1:20, and 1.0 µl was used as a template for a second round of amplification with 1.0 µM of the same degenerate primers using the Platinum Hot Start PCR Master Mix (Invitrogen). Thermocycler conditions were 94°C for 5 min; 43 cycles of 94°C for 15 seconds, 50°C for 30 sec, 68°C for 1 min.

PCR products of predicted size were excised from a 0.7% agarose/TAE gel and purified using the MinElute Gel Extraction Kit (Qiagen). Purified PCR product was ligated in the pGEM-T Easy vector (Promega) and transformed into chemically competent JM109 cells (Promega). Plasmids were purified using the QIAprep Spin Miniprep Kit (Qiagen). Sanger sequencing was performed by Retrogen (San Diego, CA).

5’ and 3’ cDNA ends were subsequently amplified by RACE PCR using the SMARTer RACE cDNA Amplification Kit and Advantage HF2 DNA polymerase (Clontech) with gene-specific primers derived from the initial RT-PCR product (Supplemental Information, Table S2). Reactions containing only the gene specific primer or the universal adaptor primer were included as negative controls. Minimum amplicon sizes were estimated using the primer positions within the full-length *X. laevis* AHR1β amino acid sequence. Amplicons exceeding the estimated minimum were gel purified as described above and cloned into the pCR Blunt II-TOPO vector using the Zero Blunt TOPO PCR Cloning Kit (Invitrogen). Ligated plasmids were transformed into One Shot TOP10 Chemically Competent *E. coli* (Invitrogen) and subsequently isolated using the QIAprep Miniprep Kit (Qiagen). Plasmids were sequenced by Retrogen (San Diego, CA). Sequences longer than 2000 bps were sequenced by University of Maine DNA Sequencing Facility (Orono, ME) using Primer Walking.

A single contiguous sequence was assembled from all RT-PCR and RACE PCR products in MacVector 14.5.3 Assembler module using the Phrap algorithm.

### Phylogenetic Analysis

Vertebrate AHR amino acid sequences were accessed from NCBI GenBank (Table S3) and aligned with the *G. multiplicata* AHR using ClustalX2. A phylogenetic tree was constructed using the well-conserved N-terminal half of each sequence by the Neighbor-Joining method and visualized in NJ-plot with 1000 bootstrap samplings. The tree was rooted with mouse ARNT as the outgroup.

### AHR expression constructs

The open reading frame encoding *G. multiplicata* AHR was synthesized by Epoch Life Sciences, Inc. with XhoI and NotI restriction sites at the 5’ and 3’ ends, respectively. The sequence was subcloned into the pCMVTNT expression plasmid (Promega). Other plasmids used in this study express *X. laevis* AHR1β (Lavine et al., 2005) and *A. mexicanum* AHR (Shoots et al., 2015), both in pCMVTNT, mouse AHR^b-1^ (provided by Dr. C.A. Bradfield, University of Wisconsin), and *X. laevis* ARNT1 (Open Biosystems, Huntsville, AL), both in pSPORT. Plasmids were transformed into JM109 chemically competent cells and grown overnight in 250 mL of LB broth with 100 µg/mL of ampicillin and purified using the HiSpeed Plasmid Maxi Kit (Qiagen).

### Velocity Sedimentation Analysis

TCDD binding was measured with a sucrose density gradient velocity sedimentation assay as described previously (Odio et al., 2013; Shoots et al., 2015). AHR constructs, expressed using the T_N_T Quick Coupled Transcription/Translation Kit (Promega), diluted 1:1 in MEDMDG buffer (25 nM MOPS, pH 7.5, 1 mM EDTA, 5 mM EGTA, 20 mM Na_2_MoO_4_, 0.02% NaN_3_, 10% glycerol, 1 mM DTT), and incubated 18 h at 4° C with 2 nM [1,6-^3^H]TCDD (Chemsyn Science Laboratories; nominal specific activity 33.1 Ci/mmol). Proteins were separated on a 10-30% sucrose density gradient by centrifugation at 60,000 rpm for 130 min in a Beckman VTi65.2 rotor. Gradients were fractionated, and the radiation content (dpm) of each fraction measured in a Beckman LS6500TD liquid scintillation counter. Unprogrammed lysate (UPL), containing the pCMVTNT empty vector, served to measure non-specific binding. Specific binding to TCDD was calculated from the radiation content in the TCDD binding peak (fractions 10-20), subtracting the background radioactivity (dpm in the UPL) in the corresponding fractions. ^14^C-labeled catalase was added to the gradient and measured as an internal sedimentation marker to allow alignment of fractions of different gradients (Supplemental Figure S1).

Relative quantity of each AHR in the assay was assessed by western blotting. Following SDS-PAGE, samples were blotted onto nitrocellulose. Blots were probed with a 1:1000 dilution of a monoclonal antibody SA210 (Enzo Life Sciences; 1 mg/mL), directed against the N-terminal half of mouse AHR and the polyclonal goat anti-rabbit secondary antibody cross-linked to alkaline phosphatase (Sigma-Aldrich). The blot was developed using the AP Conjugate Substrate Kit (Bio-Rad).

### Transactivation Assay

COS-7 cells were maintained in Dulbecco’s Modified Eagle Medium (DMEM) plus 10% Fetal Bovine Serum (FBS) at 37°C/ 5% CO_2_ with humidified air.

The transactivation activity of *G. multiplicata* was compared to other AHRs following TCDD and FICZ treatment using the Dual Luciferase Reporter Gene Assay (Promega) as described previously (Lavine et al., 2005; Odio et al., 2013; Shoots et al., 2015) with minor modifications. COS-7 cells were seeded in 96-well plates at a density of 12,500 cells per well. Cells were transfected using 0.5 μL of Lipofectamine 2000 (Invitrogen). 24 hr after plating, each well received up to five different plasmids. All reactions contained 13.4 ng of the AHR-responsive pGudLuc6.1 luciferase reporter plasmid and 1.0 ng of pTK *Renilla* (Promega). The pGudLuc6.1 plasmid uses the AHR-responsive domain from the upstream region of the mouse CYP1A1 gene to drive firefly luciferase expression (Han, Nagy, & Denison, 2004). Cells transfected with only these plasmids served to normalize for background luminescence. All other cells were co-transfected with 33.4 ng of *X. laevis* ARNT1 plasmid (Open Biosystems, Huntsville, AL) and an optimized quantity of AHR expression construct: 33.4 ng of *X. laevis* AHR1β, *A. mexicanum* AHR, or mouse AHR, or 133.6 ng of *G. multiplicata* AHR. The pCMVTNT vector, lacking an insert, was added to maintain the total amount of DNA at 200 ng per well. A negative control lacked an AHR expression vector, substituting pCMVTNT. 5 h after transfection, cells were exposed for 18 h to graded concentrations of TCDD, FICZ, or DMSO vehicle, in triplicate. DMSO concentration was held constant at 0.25% in all wells. Luminescence was measured on the GloMax Navigator (Promega) using the Dual Luciferase Reporter Assay Kit (Promega). Due to its relatively rapid metabolism, the FICZ response was also measured after 3 h exposure. In these experiments, FICZ was added after a 20 h transfection incubation, in order to maintain a 23 h total incubation time with the transfection reagent. Transcriptional response of each transfected cell culture was expressed as relative luciferase units (RLU), representing the ratio of induced firefly luciferase luminescence to constitutively expressed *Renilla* luciferase, thus normalizing for transfection efficiency. Fractional induction (Poland & Glover, 1975) of reporter gene expression was calculated by subtracting the relative luminescence of DMSO-treated cells from each value and determining its ratio the maximal response level in each concentration−response experiment. Using GraphPad Prism v.6.0h, the mean and standard error for the fractional response at each concentration were calculated and analyzed by nonlinear regression to determine the EC_50_ value for each AHR constraining the background response to 0 and the maximal response to 1.

### Homology Modeling

The structural model of *G. multiplicata* AHR ligand-binding domain (LBD; residues 281−387) was generated as described for mouse (Motto, Bordogna, Soshilov, Denison, & Bonati, 2011). Three X-ray structures of the PAS B domain of the Hypoxia-inducible factor 2α (HIF2α) in complex with artificial ligands (PDB entries: 3F1O, 3H7W, 3H82) were used as templates in MODELLER (Fiser, Do, & Sali, 2000; Marti-Renom et al., 2000; Sali & Blundell, 1993). The optimal model among the 100 generated was selected on the basis of the best DOPE SCORE (Shen, Li, & Chen, 2006). For comparison, models of the AhR LBDs of *A. mexicanum* (AmAhR; Shoots et al., 2015), *X. laevis 1b* (XlAhR; Odio et al., 2013), *G. gallus* (GgAhR; Fraccalvieri et al., 2013), which we previously developed, were generated from scratch using the same ligand-bound HIF2α complexes used for *M. musculus* (MmAhR; Motto et al., 2011) and *G. multiplicata* AHR (this work). The quality of the models was evaluated using PROCHECK (Laskowski, Macarthur, Moss, & Thornton, 1993) and the ProSA validation method (Sippl, 1993). Secondary structures were attributed by DSSPcont (Andersen, Palmer, Brunak, & Rost, 2002). The binding cavity within the modeled LBDs was characterized using the CASTp server (Dundas et al., 2006). Visualization of the models was accomplished using PYMOL (PyMol).

## Results

### Sequence analysis of *G. multiplicata* AHR

We determined the cDNA sequence of an AHR from *Gymnopis multiplicata* (the Varagua caecilian) using RT-PCR with degenerate primers and RACE PCR. The corresponding 3,742 nt mRNA contains an open reading frame of 2,532 nt, encoding an 843 residue polypeptide with predicted molecular mass of 92.7 kDa. The sequence is publicly available in the Genbank database with accession number MH457176. Sharing 59-66% identities with a wide range of tetrapod AHRs, the amino acid sequence is orthologous to all other reported amphibian AHRs (Fig. 1), including members of the orders anura (frogs and toads) and caudata (salamanders). Our phylogenetic analysis places the *Gymnopis* AHR in a monophyletic group that includes the human *AHR* as well as the *AHR* and *AHR1* genes from other vertebrate taxa. To date, no *AHR2*s have been detected in amphibians, including the well-annotated genomes of *X. tropicalis* and *X. laevis* (Lavine et al., 2005; Shoots et al., 2015) or the *Ambystoma mexicanum* transcriptome (v 4.0). To our knowledge, this cDNA represents the first reported AHR sequence from the Apoda (Gymnophiona), the first of the three extant orders to diverge from the common amphibian lineage (Pardo, Small, & Huttenlocker, 2017; Pyron & Wiens, 2011; San Mauro et al., 2014).

**Figure 1.**
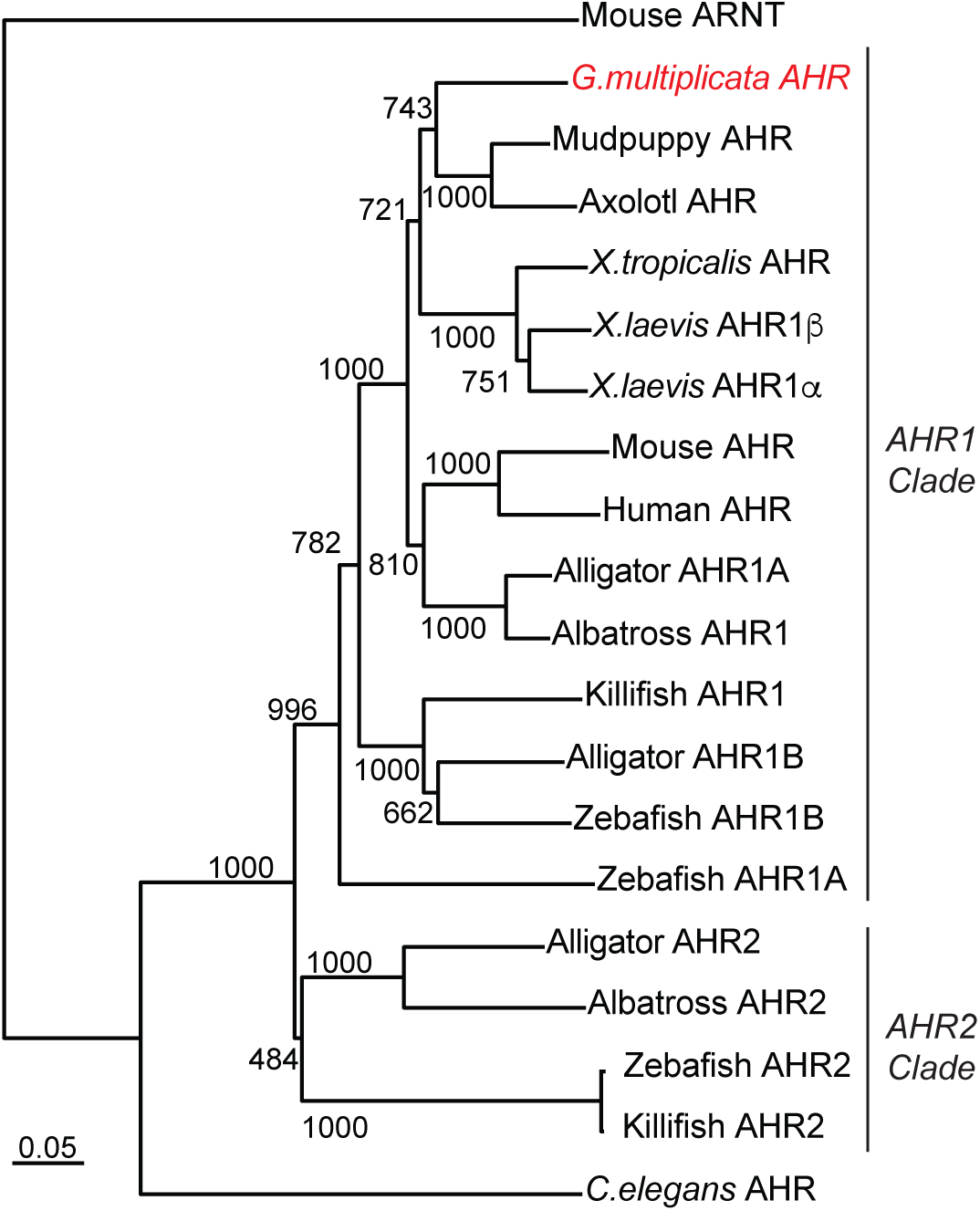
Phylogenetic analysis of *G. multiplicata* AHR. Amino acid sequences of bHLH and PAS domains of each AHR sequence were aligned in Clustal X2. A tree was inferred by the Neighbor-Joining method. Numbers at the branch points represent the bootstrap values based on 1000 samplings. Scale bar indicates degree of substitution (5 residues per 100). Accession numbers of the sequences and generic names of the species are found in Supplemental Information, Table S3.

### TCDD binding by *G. multiplicata* AHR

We sought to characterize the ligand-binding properties of the *Gymnopis* AHR, testing the hypothesis that the caecilian receptor shares the weak TCDD binding phenotype displayed by AHRs representing the Anura and Caudata (Lavine et al., 2005; Odio et al., 2013; Shoots et al., 2015). Binding to [^3^H]TCDD was measured in velocity sedimentation assays, using 2 nM TCDD, a saturating concentration for high-affinity mouse AHR^b-1^ but well below the K_d_ of *X. laevis* AHRs (Lavine et al., 2005). Equivalent quantities of all amphibian AHRs demonstrated much lower TCDD binding than the mouse receptor (Fig. 2). Consistent with our hypothesis, *Gymnopis* AHR exhibited only about one-third of the TCDD binding of mouse AHR. Surprisingly, however, it bound over 4-fold more TCDD than *X. laevis* AHR1β (frog) and *Ambystoma mexicanum* (salamander) AHRs, suggesting it binds TCDD with greater affinity than the frog and salamander receptors.

**Figure 2.**
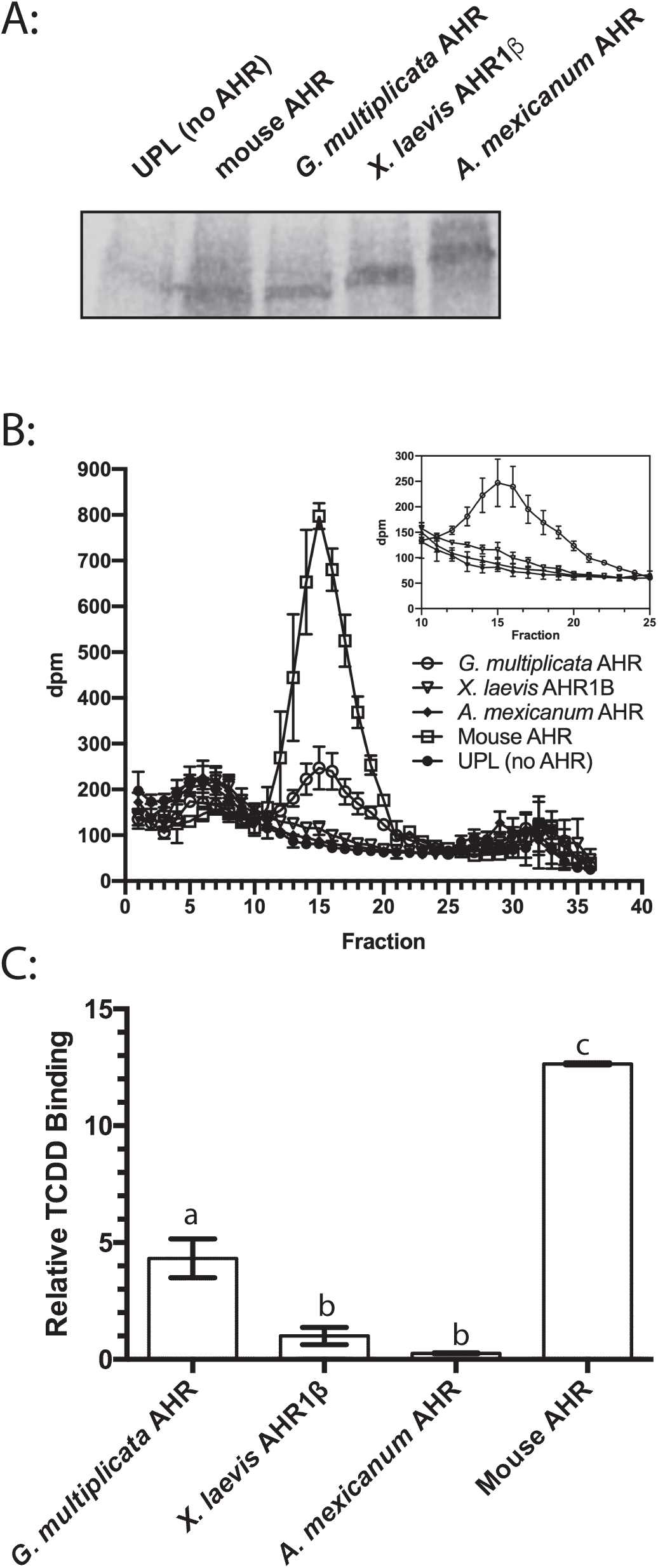
TCDD binding by *G. multiplicata* AHR. **(A)** Expression of each AHR in TNT reactions determined by Western blotting. **(B)** Velocity sedimentation analysis of TCDD binding. Synthetic AHR proteins or unprogrammed TNT lysates were incubated with 2 nM [^3^H]TCDD and fractionated on sucrose density gradients. Inset graph excludes mouse AHR and contains re-scaled y-axis. **(C)** Quantification of TCDD-specific binding revealed by sedimentation analysis. The radioactivity (disintegrations per minute) in fractions comprising each peak in panel B was summed. Specific binding is the difference between total binding (preparations containing an AHR) and nonspecific binding (preparation lacking AHR). The bar graph plots specific binding relative to that found for *X. laevis* AHR1β. Values represent means ± the standard error for four replicates. Values with identical labels do not differ statistically (One-way ANOVA with Tukey’s test for individual contrasts).

### Transactivation properties of *G. multiplicata* AHR

We next sought to determine whether the apparent weak TCDD binding properties of the caecilian AHR are reflected in its function as a ligand-activated transcription factor. These experiments made use of pGudLuc6.1, the widely-employed luciferase reporter construct driven by a strong *CYP1A1* enhancer region containing four functional dioxin responsive elements (DREs; Han et al., 2004). This reporter construct was used in previous characterizations of AHRs from *Xenopus* (Lavine et al., 2005; Odio et al., 2013) and *Ambystoma* (Shoots et al., 2015). TCDD proved to be a much less potent agonist for amphibian AHRs than for the mouse receptor (Fig. 3). While mouse AHR exhibited an EC50 in the sub-nanomolar range, values for all three amphibian AHRs were two orders of magnitude greater (Table 1). Roughly consistent with the ligand-binding assay (Fig. 2), the caecilian AHR was more sensitive to TCDD than the frog and amphibian receptors. The striking difference in efficacies (magnitude of luciferase induction) is not unusual for this assay (*e.g.*, Lavine et al., 2005; Odio et al., 2013; Shoots et al., 2015). Likely representing an artifact of *in vitro* expression level, efficacy is independent from the potency indicated by the EC50.

**Figure 3.**
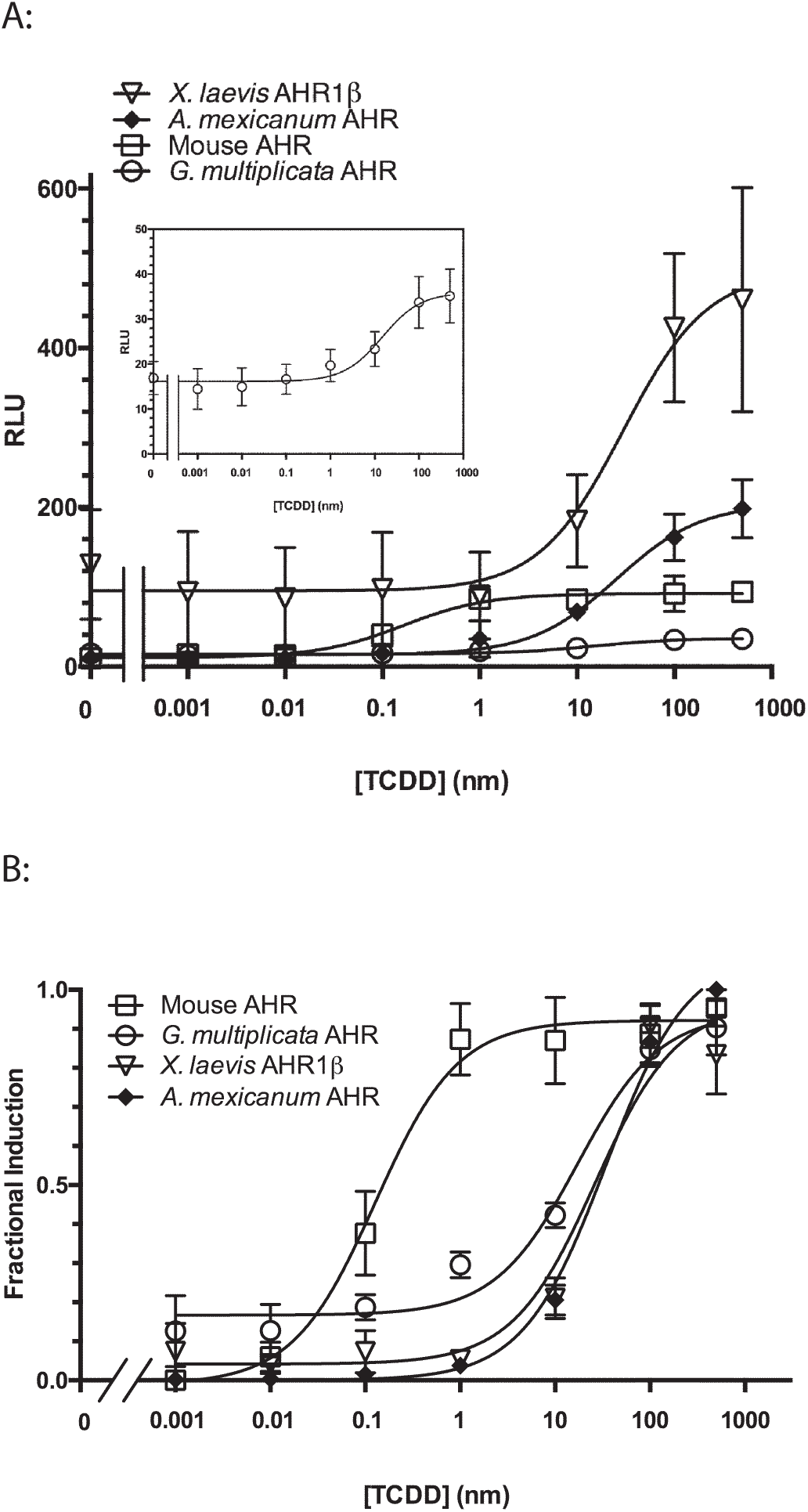
TCDD-induced transactivation activity of *G. multiplicata* AHR. COS-7 cells were cotransfected with pGudLuc6.1 reporter construct, pRL-TK transfection control construct, and expression plasmids for *X. laevis* ARNT1 and each indicated AHR. Cells were treated with DMSO or TCDD for 18 h. Each plotted value represents the mean of at least three replicate assays ± standard error. **(A)** Transactivation activity of each AHR is given in relative luciferase units (RLU), the ratio of firefly to Renilla luciferase activity at each concentration of TCDD. Inset graph depicts *G. multiplicata* AHR with re-scaled y-axis. **(B)** Fractional induction. For each AHR, relative luciferase expression at each TCDD concentration was normalized to the maximal response, which was assigned a value of 1. Nonlinear regression was used to calculate EC_50_ values for each AHR. r^2^ values for the fitted curves are 0.81 for *G. multiplicata* AHR, 0.99 for *A. mexicanum* AHR, 0.86 for frog AHR1β, and 0.92 for the mouse AHR.

**Table 1.** EC_50_ values for reporter gene induction by TCDD. COS-7 cells were transfected to express the indicated AHR and exposed to graded concentrations of TCDD for 18 h. EC_50_ values were calculated from the nonlinear regression of fractional induction against the logarithm of TCDD concentration (Fig. 3). Values represent the mean ± SE of at least four replicates.

| AHR | EC<sub>50</sub> |
| --- | --- |
| *G. multiplicata* AHR | 17.17 ± 0.03 nM |
| *X. laevis* AHR1β | 26.23 ± 0.04 nM |
| *A. mexicanum* AHR | 34.09 ± 0.02 nM |
| mouse AHR | 0.13 ± 0.03 nM |

FICZ is an exceptionally potent endogenous agonist for AHRs from a wide range of vertebrates. We measured the FICZ responsiveness of each amphibian AHR in transactivation assays. Consistent with previous studies, FICZ was 1-2 orders of magnitude more potent than TCDD in 18-hr exposures (Fig. 4a; Table 2). With the caecilian and frog receptors, FICZ potency was even greater for shorter exposure times, during which its metabolism by CYP1s and phase II enzymes intrinsic to COS-7 cells is limited (Wincent et al., 2009; Fig. 4b and Table 2). All three amphibian AHRs exhibited EC50s in the sub-nanomolar range at both time points (Table 2). The higher potency of TCDD with *Gymnopis* AHR compared to the other amphibian receptors was not evident for this endogenous agonist.

**Figure 4.**
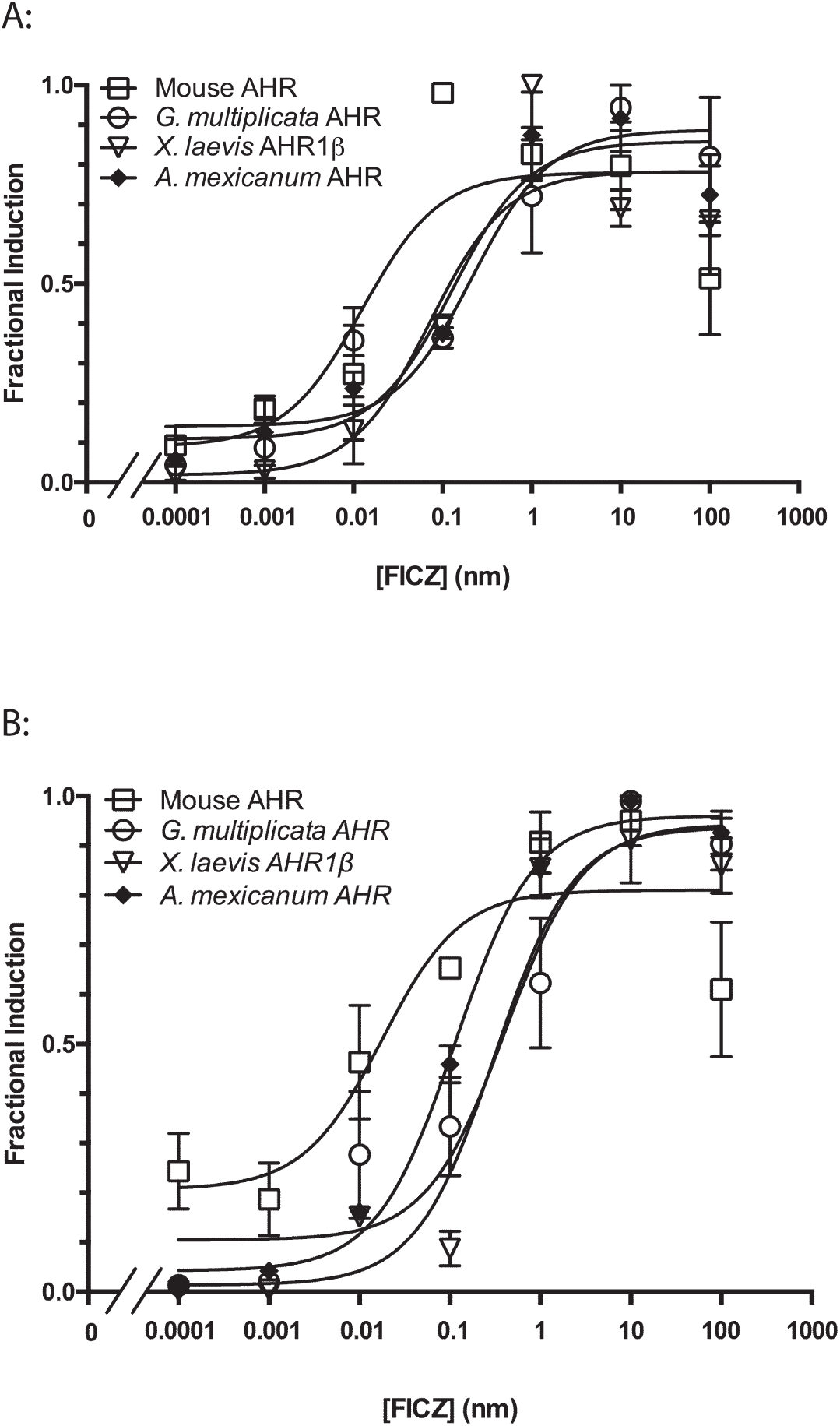
FICZ-induced transactivation activity of *G. multiplicata* AHR. Fractional induction of reporter gene expression by each AHR following **(A)** 18-h or **(B)** 3-h exposure was determined as described for Fig. 3. Values represent mean ± standard error for at least four replicates. Panel A: r^2^ values for the fitted curves are 0.86 for *G. multiplicata* AHR, 0.91 for *X. laevis* AHR1β, 0.98 for *A. mexicanum* AHR, and 0.71 for mouse AHR. Panel B: r^2^ values for the fitted curves are 0.83 for *G. multiplicata* AHR, 0.83 for *X. laevis* AHR1β, 0.85 for *A. mexicanum* AHR, and 0.65 for mouse AHR.

**Table 2.** EC_50_ values for reporter gene induction by FICZ. COS-7 cells were transfected to express the indicated AHR. EC_50_ values were calculated from the nonlinear regression of fractional induction against the logarithm of TCDD concentration. Values represent the mean ± SE of at least four replicates.

| AHR | EC<sub>50</sub> |  |
| --- | --- | --- |
|  | 18 hr | 3 hr |
| *G. multiplicata* AHR | 0.41 ± 0.04 nM | 0.21 ± 0.04 nM |
| *X. laevis* AHR1β | 0.31 ± 0.03 nM | 0.07 ± 0.04 nM |
| *A. mexicanum* AHR | 0.11 ± 0.04 nM | 0.13 ± 0.04 nM |
| mouse AHR | 0.02 ± 0.04 nM | 0.01 ± 0.04 nM |

### A structural model of the ligand-binding domain of *G. multiplicata* AHR

To explore the structural basis for *Gymnopis* AHR agonist binding and transactivation properties, we generated a homology model of the ligand binding domain (LBD; residues 281-387) using three X-ray structures of the PAS-B domain <u>of</u> Hypoxia-inducible factor 2α (HIF-2α, another sensor-type protein in the PAS family (Gu, Hogenesch, & Bradfield, 2000), in complex with artificial ligands (Motto, Bordogna, Soshilov, Denison, & Bonati, 2011; Fig. 5). An accurate structural comparison was made using the same approach to model the binding cavities of well-characterized tetrapod AHRs from frog, salamander, chicken, and mouse, which vary dramatically in their binding and responsiveness to TCDD (Fraccalvieri et al., 2013; Lavine et al., 2005; Odio et al., 2013; Shoots et al., 2015). The five 107-residue sequences shared 78 identities (72.9%; Fig. 5a). Among the 29 residues of *Gymnopis* AHR that differ in one or more species, only three bear side chains that protrude into the modeled binding cavity: N333, A362, and A378 of *Gymnopis* AHR (Fig. 5a,b). These residues are identical to the corresponding positions of *X. laevis* AHR1β and *Ambystoma* AHR that were previously demonstrated to underlie their relatively low TCDD binding (Odio et al., 2013; Shoots et al., 2015). While the high-affinity chicken and mouse receptors contain a serine in at least one of the aligned positions, the low-affinity amphibian receptors lack serine at all three (Fig. 5a). The extension of this combination of residues to a third amphibian group suggests that this LBD sequence and structural property emerged early in amphibian evolution, in the ancestor of all three extant clades, resulting in low affinity AHRs throughout the extant amphibian groups.

**Figure 5.**
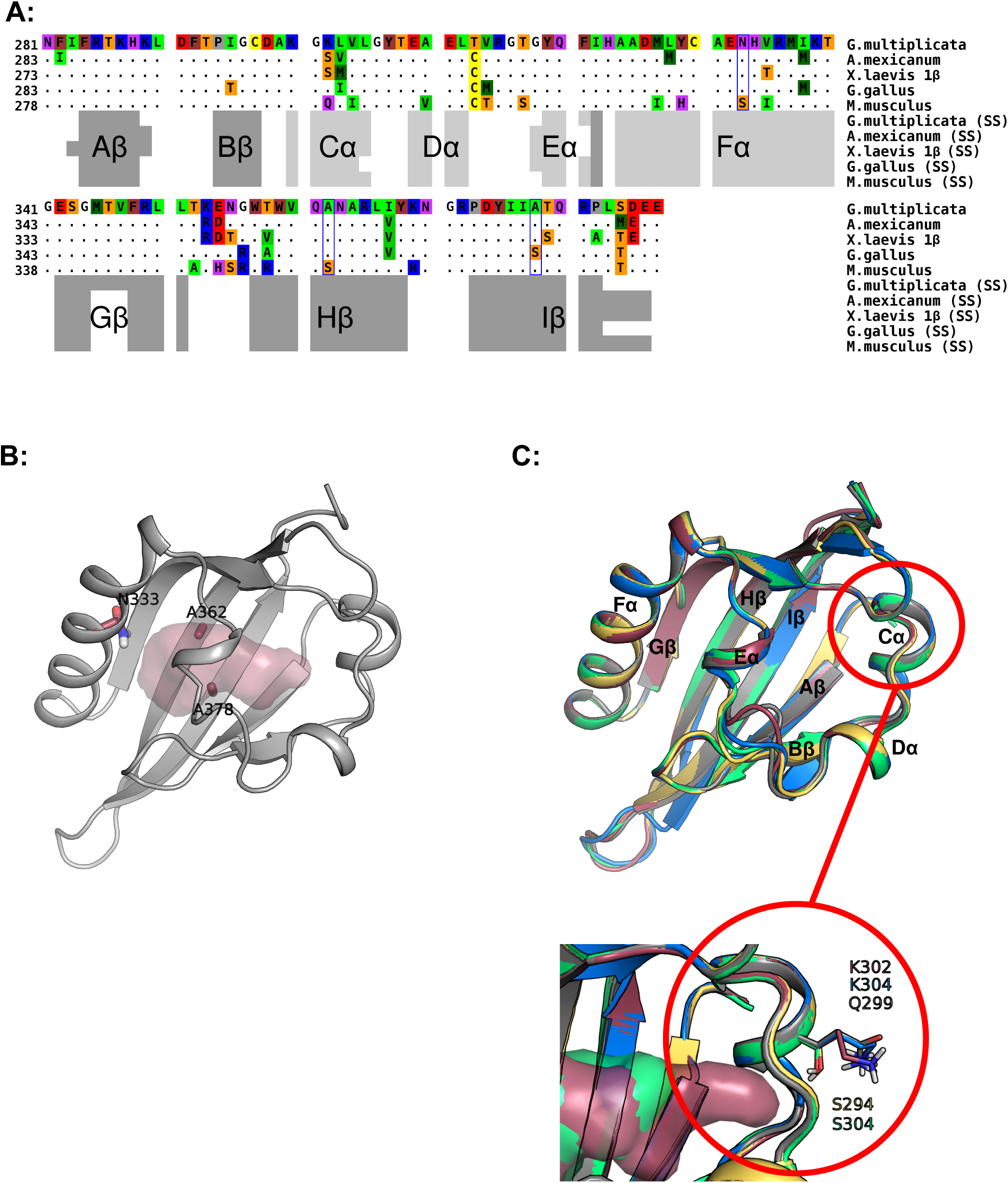
Sequence and structural model for the *G. multiplicata* AHR. **(A)** Sequence alignment produced by Clustal W. Only residues that differ from the caecilian sequence are shown; dots indicate conserved positions. Variable residues that protrude into the modeled binding cavity are boxed. Color scheme for residues: red, acidic; blue, basic; purple, polar; yellow, Cys; brown, aromatic; green, hydrophobic; orange, Ser, Thr; gray, Pro; white, Gly. Secondary structures attributed by DSSPcont (SS) are indicated below: light gray bars for helices and dark gray bars for β-strands. **(B)** Cartoon representation of the modeled *G. multiplicata* AHR LBD. Residues that both differ from the high-affinity mouse or chicken AHRs and protrude into the modeled binding cavity are labeled and shown as sticks. The light red shaded area delineates the molecular surface of the binding cavity identified by CASTp. **(C)** Comparison of cartoon renderings for modeled LBDs of AHRs: magenta for *G. multiplicata* AHR, green for A*. mexicanum* AHR, yellow for frog AHR1β, blue for chicken AHR, and gray for mouse AHR. Black labels indicate the conserved secondary structure elements identified by DSSPcont. Magnified inset image depicts position of potentially variable residues that protrude outside the binding pockets but may nonetheless affect binding cavity dimensions (see text).

## Discussion

### Conservation of low dioxin affinity in diverse Amphibian AHRs

Previous characterizations of xenobiotic toxicity in amphibians reveal that frogs and salamanders are generally insensitive to toxicity of dioxin-like chemicals (Beatty et al., 1976; Collier et al., 2008; Jung & Walker, 1997; Pezdirc et al., 2011; Vajda & Norris, 2005). Low-affinity agonist binding by their AHRs is likely a major mechanistic contributor to this phenotype (Lavine et al., 2005; Odio et al., 2013; Shoots et al., 2015). In this regard, these animals differ from both teleosts, which diverged from the vertebrate lineage prior to amphibians, and the other tetrapod groups, which arose thereafter. Thus, it seems likely that high-affinity dioxin binding by AHR was lost in a lineage-specific fashion. In frogs (Order Anura) and salamanders (Urodela), low-affinity dioxin binding is conferred largely by the lack of a serine residue in at least one of three key positions within the modeled ligand-binding cavity (Odio et al., 2013; Shoots et al., 2015). The present study reveals that the same absence of serines at the aligned binding cavity positions within the AHR sequence isolated from the Varagua caecilian (*Gymnopis multiplicata*; Apoda). The encoded protein also exhibits relatively low dioxin sensitivity. This finding is consistent with the hypothesis that low dioxin affinity emerged early in amphibian evolution, in a common ancestor of all three extant orders.

While the asparagine and the two alanines associated with low TCDD affinity in the three previously characterized amphibian AHRs are well conserved, the TCDD binding and sensitivity of the caecilian receptor are measurably greater (Figs. 2, 3). This suggests that additional, more subtle structural features may also be influential. One such feature is the overall volume of the modeled binding cavity. Our estimation of the cavity volume in the models of all five proteins suggests that the caecilian AHR contains a larger cavity than its amphibian orthologs: 675 Å^3^, compared with 517 Å^3^ for *Ambystoma* AHR and 595 Å^3^ for *X. laevis* AHR1β. The high-affinity AHRs have even larger modeled cavities: 802 Å^3^ for mouse AHR and 715 Å^3^ for the chicken receptor. This seeming correlation suggests the hypothesis that larger cavity volume enables tighter TCDD binding. Possibly underlying the differences in cavity volume is a variable residue in the helical Cα region (Figure 5c). There is only one non-conservative substitution in the LBD of these five AHRs. This position contains serine in the small-cavity AHRs of frog (S294) and salamander (S304), corresponding to glutamine in the mouse receptor (Q299) and lysine in the caecilian (K302) and chicken (K304) proteins. We speculate that the longer, charged side chains may tend toward the solvent, leaving the cavity wider in the Cα/Dα region, while the shorter, polar serines may interact with internal residues, leading to a more compact cavity. There is reason to view this hypothesis with caution. Cavity volume and ligand binding typically relate to the conformations of amino acid side chains that protrude inward, exact conformations of which cannot be predicted with high accuracy by homology modeling. Future studies will use mutagenesis to directly test hypotheses about the relationship between cavity volume, internal side chains, and external side chains in the binding of TCDD and other agonists.

FICZ, an endogenous AHR agonist, exhibits a much smaller degree of interspecies difference than TCDD in potency for reporter gene activation by amphibian and mouse receptors, ranging over 260-fold for TCDD, but only 21-fold for FICZ (Fig. 4; Table 2). This compound engages the LBD through a different set of interactions than the dioxin-like compounds, as evidenced by the distinct effects of amino acid changes on receptor activation by these different ligands (Soshilov & Denison, 2014). The relative conservation of the endogenous compound’s high potency suggests that relatively robust FICZ binding—and the related non-toxicological effects—may have been maintained under stronger selective pressure than dioxin binding and subsequent receptor activation. Under this hypothesis, dioxin binding could represent one of AHR’s pleiotropic functions that has been diminished in amphibians following stochastic sequence change in an early common ancestor, even while FICZ binding and responsiveness have persisted in stronger fashion. Alternatively, the loss of AHR sequence elements associated with high TCDD potency may have been adaptive in early amphibians despite continued selection for effective FICZ agonism. In either case, the greater potency of FICZ relative to dioxin-like compounds seems to have emerged in a common ancestor and to have been conserved in all three modern amphibian groups.

### Loss of multiple AHR paralogs in the Amphibian lineage

Amphibians AHRs differ from those in the other vertebrate groups in a second important respect—gene number. AHR-related genes can be sub-divided into five different categories in the jawed vertebrates: four distinct AHR genes (AHR, AHR1, AHR2, AHR3) plus the AHR Repressor (AHRR; Hahn et al., 2017). While humans and rodents have only one AHR, most vertebrates—including cartilaginous fish, teleosts, lobe-finned fish, reptiles, birds, and many mammals—harbor members of at least two of the four AHR gene types and in some cases multiple paralogs of each (Hahn et al., 2017). In the amphibians for which AHRs have been sought by molecular cloning and/or genome searches, a single AHR seems typical. Only *X. laevis* harbored more than one AHR gene (Lavine et al., 2005), the result of a relatively recent genome duplication that arose as part of an interspecies hybridization between ancestral *Xenopus* species (Session et al., 2016). Using degenerate primers that successfully targeted multiple AHRs in other vertebrate groups, we detected only one cDNA sequence in *G. multiplicata* (this study) and *Ambystoma mexicanum* (Shoots et al., 2015), and our searches of the *Ambystoma* genome and transcriptome did not reveal the existence of additional AHR paralogs. Thus, we suggest that the presence of only one AHR may reflect the loss of multiple paralogs in an early amphibian ancestor, perhaps coincident with the loss of high-affinity agonist binding. These changes may have resulted predominantly from unknown selective pressures related to the unique features of amphibian development and/or the increasingly terrestrial lifestyle that emerged in early amphibian evolution. Alternatively, the loss of AHR genes could derive from genetic drift and a subsequent bottleneck soon after the amphibian divergence from the common vertebrate lineage.

Phylogenetic analysis (including Fig. 1) places *G. multiplicata* and all other amphibian AHRs in a monophyletic group, part of a larger clade that includes mammalian AHRs and AHR1s from other vertebrates. This result is consistent with the AHR relationships reported in many publications spanning two decades (*e.g.*, Hahn et al., 2006; Hahn, Karchner, Shapiro, & Perera, 1997; Shoots et al., 2015). A more recently published analysis, which considers conserved synteny, along with the identification of two distinct “AHR1” forms in some species including chicken and alligator (Lee et al., 2013; Oka et al., 2016), offer the possibility that AHR (first identified in humans and rodents) and AHR1 (initially identified in teleosts) are not orthologs but rather representatives of distinct paralogous groups (Hahn et al., 2017). These analyses identify the *Xenopus* genes as “*AHR*s,” not “*AHR1*s,” as their original names suggest. The close relationship of the salamander and caecilian receptors suggests that they may also be *AHR*s. Confirmation of these relationships by synteny analysis awaits a more thorough mapping and annotation of the *Ambystoma* genome and the development of an annotated, searchable caecilian genome sequence. Isolation of additional *AHR* gene and cDNA sequences from other amphibian species will also be important for this determination.

### Conclusion

Several reports have pointed out that the unique position of Order Apoda makes it important in the study of amphibian evolution (*e.g*., Case & Wake, 1975; Wu, Liu, Meng, Zhang, & Liang, 2015; Zhang & Wake, 2009). This class was the first to diverge from the common amphibian lineage after the split from the rest of the vertebrates. Furthermore, analysis of the Hox genes of a Banna caecilian (*Ichthyophis bannanicus*) found that members of this group have a relatively slow evolution rate (Wu et al., 2015). Their early divergence and their slow sequence change rate combine to suggest that caecilians can provide key insights into the traits of ancestral amphibians.

This characterization of the *G. mutliplicata* AHR has important implications for our understanding of the evolution of amphibian AHRs. This caecilian AHR shares several characteristics that distinguish other amphibian receptors, including sequence, structure, low TCDD binding affinity, and low TCDD responsiveness. This suggests that the low-affinity phenotype arose after the divergence of amphibians from the common vertebrate lineage, but prior to the divergence of the Apoda order from the other amphibians. However, only four amphibian AHRs have been extensively characterized thus far (*X. laevis* AHR1α and AHR1*β, A. mexicanum* AHR, and *G. multiplicata* AHR). A larger, phylogenetically diverse sampling of amphibian AHRs should be characterized in order to more fully test this hypothesis.

## Acknowledgements

We thank the Museum of Vertebrate Zoology at the University of California, Berkeley (Dr. Carol A. Spencer, Staff Curator of Herpetology) for providing the tissue sample from *Gymnopis multiplicata*. This work was funded by the National Institute of Environmental Health Sciences: ES011130 (WHP), ES07685 (MSD), ES006272 (MEH). Additional funding was from the Kenyon College Summer Science Scholars program.

## Conflict of Interest

Authors have no conflicting interests to declare.

## Data Sharing

cDNA sequence is publicly available in the Genbank database with accession number MH457176. Other data that support the findings of this study are available from the corresponding author upon reasonable request.

