## Supplemental Information for "An Aryl Hydrocarbon Receptor from the Caecilian *Gymnopis multiplicata* Suggests Low Dioxin Affinity in the Ancestor of All Three Amphibian Orders"

Contents:

Table S1: Degenerate primers for RT-PCR
Table S2: Gene-specific RACE primers
Table S3. Accession numbers of sequences used in phylogenetic analysis.
Figure S1. $^{14}C$-catalase marker fractionation.

**Table S1. Degenerate primers for RT-PCR.** Degenerate primers were designed from conserved regions of amino acid sequence of several vertebrate AHRs (Hahn & Karchner, 1995).

| Name | Sequence |
|---|---|
| Qf | 5'-AACCCITCIAAGMGICAYMG-3' |
| A1 | 5'-CGGGATCCARGCICTSAAYGGITT-3' |
| A2 | 5'-CGGGATCCGAYTAYCTIGGITTYCARCA-3' |
| B1 | 5'-GCTCTAGACATICCRCTYTCICCIGTYTT-3' |
| B2 | 5'-GCTCTAGAGCTCIRCYTCIGTRTAICC-3' |

**Table S2. Gene-specific RACE primers.**

| Primer Name | Product | Sequence |
|---|---|---|
| 5' Mult R | 5' RACE | 5'-CCTGGGAAAAGCCATCATCTCCTTGAG-3' |
| 3'GSP_330 | 3' RACE | 5'-AGAGGGAACCGTTGAGGCTGCTG-3' |
| 3'GSP_510 | 3' RACE | 5'-GTCACTTGGGGATTCCACCTCGC-3' |
| 3'GSP_696 | 3' RACE | 5'-GTCGCCTTTGCTGGAGGAAAACAAGATG-3' |

**Table S3. Accession numbers of sequences used in phylogenetic analysis.**

| Common Name | Generic Name | Protein | Accession Number |
|---|---|---|---|
| Varagua caecilian | *Gymnopis multiplicata* | AHR | MH457176 |
| Mexican axolotl | *Ambystoma mexicanum* | AHR | KM660691 |
| Mudpuppy | *Necturus maculosus* | AHR | AAF15278 |
| Western clawed frog | *Xenopus tropicalis* | AHR | XP002933348 |
| African clawed frog | *Xenopus laevis* | AHR1α | NP_001165383 |
| | | AHR1β | AA V49748 |
| Mouse | *Mus muculus* | AHR | NP_038492 |
| Human | *Homo sapiens* | AHR | L19872 |
| Black-footed Albatross | *Phoebastria nigripes* | AHR1 | BAC87795 |
| | | AHR2 | BAC87796 |
| American alligator | *Alligator mississippiensis* | AHR1A | BAV14369.1 |
| | | AHR1B | BAV14370.1 |
| | | AHR2 | BAV14371.1 |
| Zebrafish | *Danio rerio* | AHR1B | NP001019987 |
| | | AHR1A | AAM08127 |
| | | AHR2 | U29679 |
| Killifish | *Fundulus heteroclitus* | AHR1 | AAC60334 |
| | | AHR2 | AAC59696 |
| Nematode | *Caenorhaditis elegans* | AHR | AAC00000 |
| Mouse | *Mus musculus* | ARNT | AAA56717 |

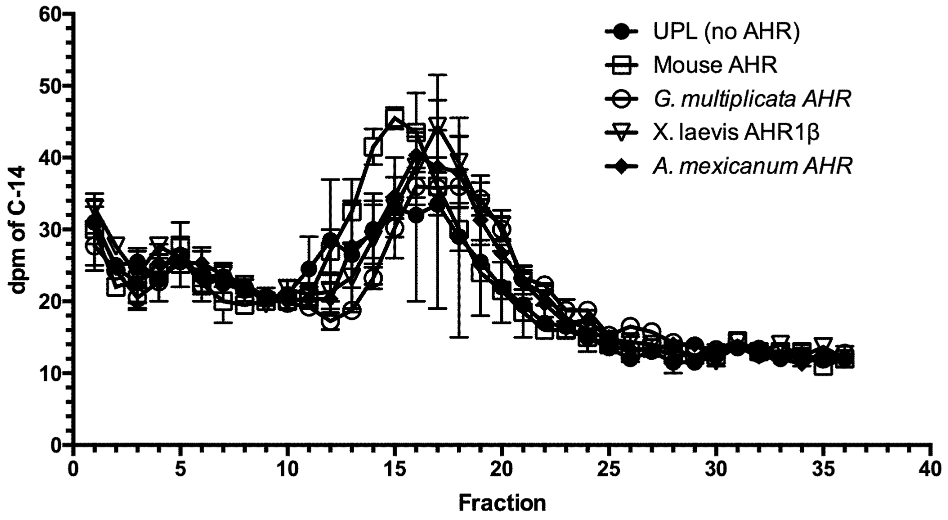


**Figure S1. $^{14}$C-catalase marker fractionation**. $^{14}$C-catalase standard (11.3S) was added to all sucrose density gradients in addition to the AHR proteins and $^{3}$H-TCDD. Gradients were fractionated and the radioactivity (dpm) was quantified over 15 minutes for each fraction by liquid scintillation counting. This standard served as a marker that could capture variability in protein mobility on the gradients and enable alignment of fractions from different gradients, since differences in elution position mirror the differences in $^{3}$H-TCDD peaks corresponding to specific binding of AHR. n=4 for each AHR.
